# A new adaptive procedure for estimating perceptual thresholds: the effects of observer bias and its correction

**DOI:** 10.1101/2021.06.15.448359

**Authors:** Chiara Fioravanti, Christoph Braun, Axel Lindner, Sergio Ruiz, Ranganatha Sitaram, Diljit Singh Kajal

**Author notes:** Corresponding author: Dr. Diljit Singh Kajal, Department of Neurology, Universitätsklinikum Frankfurt, Brain Imaging Center, Schleusenweg 2-16, 60528 Frankfurt, Germany.

## Abstract

Adaptive threshold estimation procedures sample close to a subject’s perceptual threshold by dynamically adapting the stimulation based on the subject’s performance. Yet, perceptual thresholds not only depend on the observers’ sensory capabilities but also on any bias in terms of their expectations and response preferences, thus distorting the precision of the threshold estimates. Using the framework of signal detection theory (SDT), independent estimates of both, an observer’s sensitivity and internal processing bias can be delineated from threshold estimates. While this approach is commonly available for estimation procedures engaging the method of constant stimuli (MCS), correction procedures for adaptive methods (AM) are only scarcely applied. In this article, we introduce a new AM that takes individual biases into account, and that allows for a bias-corrected assessment of subjects’ sensitivity. This novel AM is validated with simulations and compared to a typical MCS-procedure, for which the implementation of bias correction has been previously demonstrated.

Comparing AM and MCS demonstrates the viability of the presented AM. Besides its feasibility, the results of the simulation reveal both, advantages, and limitations of the proposed AM. The procedure has considerable practical implications, in particular for the design of shaping procedures in sensory training experiments, in which task difficulty has to be constantly adapted to an observer’s performance, to improve training efficiency.

## Introduction

Perceptual thresholds might vary due to different variables such as fatigue, fluctuations of attention, or sensory learning (Gorea & Sagi, 2000). Adaptive threshold estimation procedures are most effective by providing quasi-instantaneous estimates of an otherwise fluctuating sensory threshold. These estimates are much needed (Fechner, 1860; Swets, 1961), for example, in experiments in which the sensory stimulation should be kept close to an individual’s threshold, like in sensory learning experiments. In this type of experiment the level of challenge should be maintained throughout the task to achieve optimal learning. A common problem related to all threshold estimation procedures is that the thresholds reflect not only the individual’s sensitivity but also their internal processing biases. ‘Bias’ suggests a systemic tendency of the observers to over- or under-estimate the stimulus parameters (Macmillan & Creelman, 1990, 2004). There are several types of biases that can occur at different stages of perceptual processing: at the sensory level (e.g., due to sensory adaptation), at the decision-making level (e.g., due to a preference of one stimulation condition over another), the response selection level (e.g., a general preference to rather respond with the right than with the left hand in bimanual response tasks). Accordingly, any of the aforementioned internal processing biases can significantly distort sensory threshold estimates. The observer’s bias might be reduced by an appropriate design of the threshold detection experiment or it can be corrected during subsequent data analysis (Lynn & Barrett, 2014; McNicol, 2005; Swets, 2014). Nevertheless, there are occasions where an online bias correction is mandatory. Signal detection theory (SDT) is well established as a tool to independently assess an individual’s sensitivity and bias by modeling perception as a decision-making process (Gorea & Sagi, 2000; Green & Birdsall, 1978; Harvey Jr, 1992; Klein, 2001; Macmillan & Creelman, 1990, 2004; Macmillan, Rotello, & Miller, 2004; Swets, 1961; Wickens, 2002) [for details see supplementary material 1]. In SDT, a stimulus is thought to elicit a defined sensation which leads to the selection of one out of two responses with a certain probability. In this framework, two different stimuli, are represented by two probability density functions that are shifted depending on how differently they are perceived. In SDT, *d’* (the distance between the peaks of the two probability density functions, describing response behavior for individual stimuli) is increasing with stimulus discriminability. Based on SDT, the criterion for the sensory decision is a function of the individual’s perceptual bias and defines the probabilities for either of the alternative responses for each stimulus (Gorea & Sagi, 2000)(Fig. 2).

In fact, for some threshold estimation procedures, such as the method of constant stimuli (MCS), bias correction procedures provided by SDT are readily established (Maniscalco & Lau, 2014; Stanislaw & Todorov, 1999). However, this is not the case for the adaptive procedures, which – as compared to the mentioned procedures – have the advantage of providing quasi-instantaneous threshold estimates. Therefore, the present study introduces a new adaptive procedure that combines the advantages of adaptive threshold estimation procedures with the capability to correct the subject’s response bias (Kajal, 2018; Kajal et al., 2020).

The rationale behind the proposed approach is explained and investigated through several simulations demonstrating the feasibility of the procedure. Furthermore, the prerequisites, advantages, and limitations of the approach are discussed. To validate the new method, it is compared to a standard non-adaptive procedure, the MCS. Amongst the variants of MCS application, the “classic” version – which is distinct from AM – was chosen, as it is not involving any adaptation based on subjects’ responses. To illustrate practical application of the proposed bias-corrected adaptive threshold estimation procedure, it is simulated and discussed in the context of a visual backward masking paradigm (Del Cul, Dehaene, & Leboyer, 2006); also see (Di Lollo, Enns, & Rensink, 2000; Enns & Di Lollo, 2000; Vorberg, Mattler, Heinecke, Schmidt, & Schwarzbach, 2003).

## Methods

Testing reliability and validation of our new adaptive method for the estimation of the bias-corrected perceptual threshold were carried through various simulation procedures. To have experimentally contrived sensory capabilities and perceptual biases, we specifically designated a virtual observer (VO) within the framework of signal detection theory (SDT) (Blake, Bülthoff, & Sheinberg, 1993; Crary, 1990). In a first step, the estimation of the VO’s perceptual threshold was simulated, using the MCS and the chosen AM procedure, with and without bias correction. Using MCS and AM effects of trial number and bias strength on the observer’s threshold and bias estimates are investigated. Moreover, results obtained for comparing AM and MCS by simulating time-varying sensitivities and linearly changing biases are presented.

### Perceptual threshold estimation procedures

In the following paragraph, we briefly describe the method of constant stimuli and the adaptive method:

a. **Method of constant stimuli (MCS)** MCS refers to a procedure in which a set of preselected stimuli are presented with stimulus parameters that could cover the whole perceptual range i.e. 0% to 100% correct responses (McKee, Klein, & Teller, 1985; Treutwein, 1995). Offline estimation of the perceptual thresholds is performed by fitting a psychometric function that relates to the observer’s response pattern as a function of stimulus parameters. The sensory threshold for a given performance, i.e., 75 %, 70.7 % or 66.7 % is derived from the inverse of the psychometric function.
b. **Adaptive method (AM)** AM approximates towards stimulus parameters that lead to a predefined performance level (e.g., 70.7%, or 66.7% of correct responses depending on the adaptation rule used for stimulus selection). This is achieved by varying stimulus parameters across trials, based on the observer’s responses in the preceding trials (Treutwein, 1995; Watson & Pelli, 1983). In AMs, stimulus parameters tends to sample more densely around the individual’s perceptual threshold value (Levitt, 1971). Furthermore, AMs are regarded as being more efficient in terms of time since a smaller number of trials are needed (Watson and Fitzhugh, 1990). Furthermore, AMs can also provide quasi-instantaneous threshold estimates.

### Virtual Experiment

For the simulations of the threshold estimation procedures a virtual experiment was conducted in which a virtual observer judged the emotional valence face images. Each trial (Fig.1) of the virtual experiment started with the presentation of a prime stimulus for the time duration of 16.67 ms accommodating either an emotionally positive (happy) or negative (sad) face. After a given time delay, the prime stimulus was masked by an emotionally neutral face, of the same identity as of prime stimulus, for the time duration of 250 ms. In such a paradigm, the emotional content of the prime stimulus cannot be correctly identified for a time delay of zero between prime and mask stimuli, on the contrary the probability for correctly identifying the emotional content increases with the increase in the duration of the time delay. A black screen for the target time delay duration is displayed between the prime and the mask stimuli. The respective time delay durations could correspond to one of ten different values Δ*t* (16.67 ms x k, 0 ≤ k ≤ 9). After the presentation of the mask, a black screen was displayed. To indicate the emotional valence of the prime (negative or positive), one of the two virtual response buttons were selected by the virtual observer.

**Fig. 1:** Backward masking paradigm. In the original setup to which the simulation refers, the assignment of the response buttons was randomly altered on a trial-by-trial basis. The instruction “Neg + Pos” informed the subject that the left button should be pressed if the emotion of the prime was negative and the right button should be pressed if the emotion was positive. The “Pos + Neg” indicated the reverse assignment.

#### On request

During the simulation for the MCS approach, 10 different predefined delays were presented across trials in a randomized order (Leek, 2001). To estimate the perceptual threshold, a sigmoid psychometric function (logistic regression) was fitted to the probability of correct responses as a function of the predefined delays. The threshold delay was determined for a level of correct identification of the emotional expression of 66.7%.

For the simulation of the AMs, the ‘two-down one-up rule’(Leek, 2001) was applied to select the time delay between prime and mask stimuli in the upcoming trial. In this procedure, the time delay between the prime and the mask decreases by one step (16.67 ms) after two correct responses and increases by one step with each incorrect response. Assuming a stationary threshold, the delay can be expected to asymptotically approach the threshold. In a two-down one-up rule, in which the number of correct responses does not require to be in a consecutive sequence, the stimulation will converge to the performance level of 66.7 % of correct responses. Differently, when correct responses are requested to appear consecutively to decrease the delay by one step, the performance level would converge to 70.7 % correct responses (Leek, 2001)(see supplementary material [2]).

### A new adaptive method with bias correction procedure

Our newly proposed method for threshold estimation integrates the advantages of an adaptive method and SDT based bias-correction procedure. According to SDT, the response outcome of an observer depends on the position of the criterion *γ* in the probability density distribution that describes stimulus perception. In our method, the probability density functions are centered at 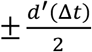. Fixed criteria *γ* were applied for sad and happy faces for a single mask delay Δ*t* (Gorea & Sagi, 2000). Assuming a differential change of the detectability for sad and happy faces across mask delays as a result of sensory bias, individual criteria for each mask delay were estimated. In the framework of the SDT, bias correction corresponds to a shift of the criterion to the common center of both Gaussian distributions. In Fig. 2, the bias-free criterion corresponds to *γ*_*c*_ *= 0* solid vertical line) and the biased criterion is *γ* (dashed vertical line). Given an estimate of the bias at *γ*, the bias correction procedure determines the observer’s response, if the criterion was at the center point of both probability density distributions.

**Fig. 2:**
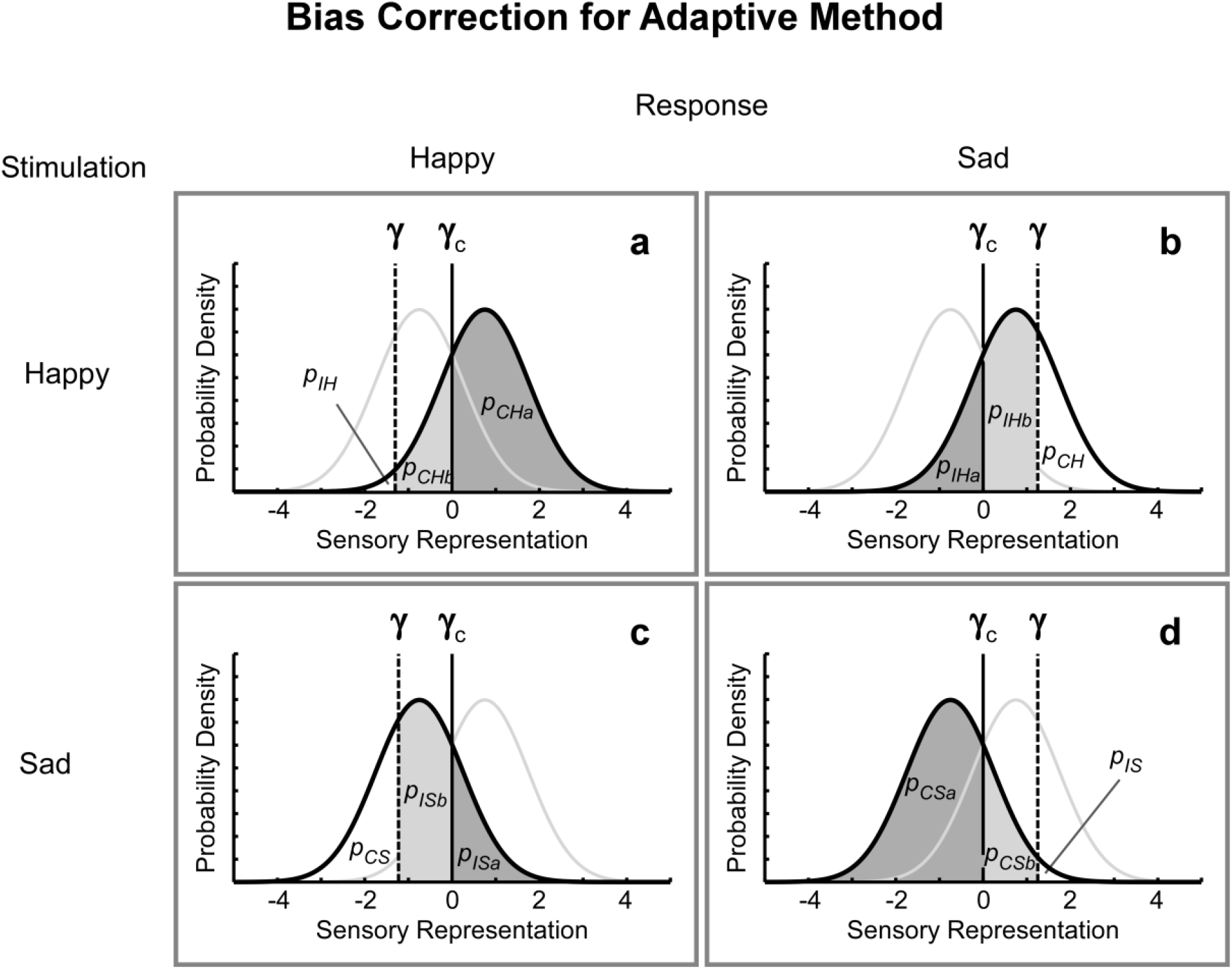
In the adaptive method with bias correction, the observer’s bias γ is estimated, and the observer’s response is corrected by eliminating any bias (γ_c_ = 0) Based on the corrected response, the stimulus for the next trial is chosen according to the two-down one-up procedure. Since in a single trial it is unknown how much the decision reflects the bias, a probabilistic correction needs to be applied. For this purpose, probabilities for correctly identified happy faces (P_CS_ (λ)), for correctly identified sad faces (P_CS_ (λ)), for incorrectly identified happy faces (P_IH_ (λ)), and for incorrectly identified sad faces (P_IS_ (λ)),will be split into a part that describes the probability for the unbiased criterion γ_c_ =0 (solid vertical line) and the proportion due to the bias-dependent criterion *γ* (dashed vertical line). The unbiased proportion of the probabilities refers to p_•a_ and the component that is due to the response bias to p_•a._ Depending on the stimulation and the response, the ‘•’symbol represents correctly or incorrectly identified happy faces or sad faces (CH, IH, CS, IS), in a) and b) HF stimuli and in c) and d) SF stimuli are presented. In a) and c) the correction needs to be considered for the response HF, and in b) and d) for the response SF.

Likewise, in the standard AMs procedures, our approach estimates the threshold using the “two-down one-up” rule. The procedure starts without any bias correction. During the experiment, the bias for the adaptive procedure is updated for each trial and will be subsequently used to determine the bias-corrected observer’s response. The bias estimate for the current trial for the respective time-delay are based on all previous trials within the framework of SDT (explained in detail in the next paragraph). The delay to be chosen for the upcoming trial is based on two prerequisites: first, valuation of a bias-corrected response (Fig.2), and second, application of the two-down one-up rule. Based on the estimated location of the criterion, the bias-corrected observer’s response might either be accept or reversed, i.e., a “negative emotion” response could be turned into a “positive emotion” response or vice versa. The algorithm determining the bias corrected response in a single trial is based on probabilistic considerations and is explained in detail in the following paragraph.

The rules for selecting the stimulus parameters for the next trial in the AM are summarized in the flowchart (Fig. 3). To define the mask delay of the following trial, four different conditions need to be considered in the approach (Fig. 2)

a. Assuming that in a certain trial a ‘happy face’ (HF) is presented and the observer’s criterion to classify the stimulus as ‘happy’ or ‘sad’ is at a level of *γ* < 0 (*this situation describes the case of a bias towards happy faces*) (Fig. 2a), the probability to choose ‘sad face’ as a response corresponds to the rate of *p*_*IS*_ (*λ*) In contrast, the probability to select HF as a response is referred to as *p*_*CH*_ (*λ*) The probability *p*_*CH*_ (*τ*) can be thought of as being composed of *p*_*CH*_ *= p*_*CHa*_ *+ p*_*CHb*_, where 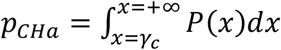 is the cumulative probability for *X > γ*_*c*_ and 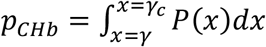 is the probability for*γ <x ≤ γ*_*c*._ *γ < 0*represents an observer’s perception criterion and *γ*_*c*._ = 0 the bias-free perception criterion. In other words, *p*_*CHa*_ refers to the cumulative probability to identify the happy face in case of no bias, and *p*_*CHb*_ to the part of the probability that is due to the bias. *P(x*) is assumed to be normally distributed. In the proposed method, the selection of the next stimulus is based on an observer’s bias-corrected response. If the virtual observer responded with a ‘sad face’ (SF), the response was wrong, regardless of any potential bias (Fig. 2). However, given the response bias towards HF, whenever the virtual observer answers HF, only a proportion 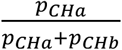 of these responses can be accepted as HF. For the remaining responses, bias-correction will be converting the decision to SF. In an individual trial, depending on the estimated proportion, the response will be kept or changed. In detail, a number *r* will be drawn from a uniform distribution between 0 and 1. If 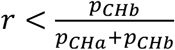, the observer’s response will be changed from HF to SF, i.e., from correct to incorrect.
b. Likewise, for HF stimuli, an HF response will remain unchanged when the criterion is set at a level of *γ > 0 (this situation describes the case of a bias towards sad faces*) (Fig. 2b). The probability of incorrectly perceiving a happy face *p*_*IH*_ (*τ*), i.e. responding with SF to the HF stimulus, can be split into a proportion depending on the bias-free criterion *p*_*IHa*_ and a proportion that corresponds to the observer’s bias *p*_*IHb*_ If it holds for the selection variable 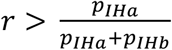, the SF response will be changed to HF.
c. For SF stimuli, an SF response will remain unchanged when the criterion is set at a level of *γ < 0 (this situation describes the case of a bias towards happy faces in the presence of an SF stimulus*) (Fig. 2c). The probability of *p*_*IS*_ (responding with HF to the SF stimulus), can be split into a proportion depending on the bias-free criterion *p*_*Isa*_ and a proportion corresponding to the observer’s bias *p*_*Isb*_. In the case of 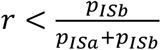 the HF response will be changed to SF.
d. Finally, for SF stimuli a HF response will remain unchanged if the criterion is set at a level of *γ > 0 (describing a bias towards sad faces*) (Fig. 2d). The probability of correctly perceiving a sad face *p*_*CS*_ i.e., responding with SF to the SF stimulus, can be split into a component depending on the bias-free criterion *p*_*CSa*_ and a proportion that corresponds to the observer’s bias *p*_*CSb*_ If it holds for the variable *r* drawn from a uniform distribution between 0 and 1,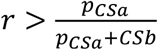, the SF response will be changed to HF. In all four cases the bias-corrected response is used for the selection of the next stimulus according to the two-down one-up rule in the subsequent step of the algorithm.

**Fig 3:**
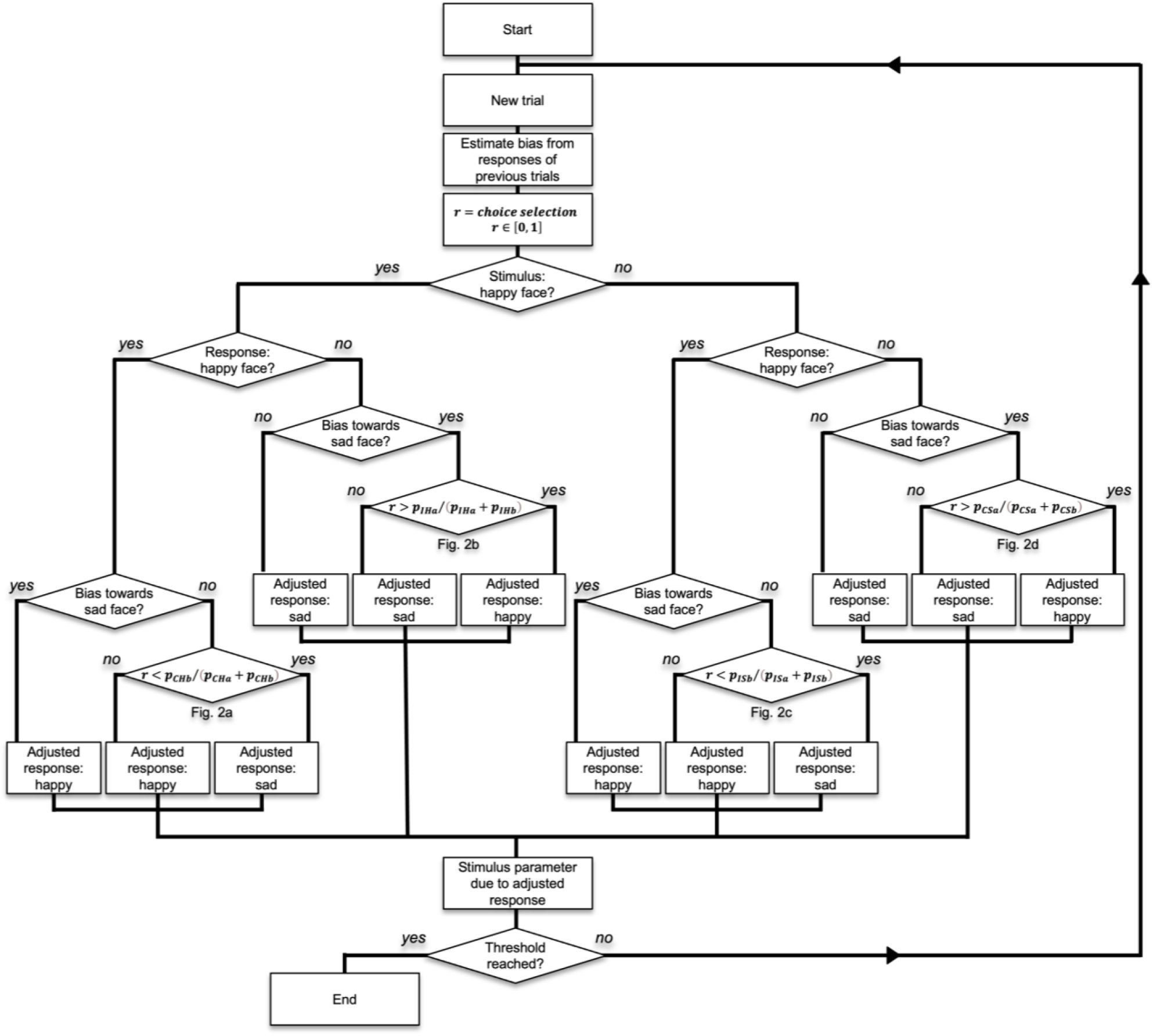
Flowchart explaining the procedure for bias correction in AM. A bias towards sad faces corresponds to a criterion *γ* >0 vice versa a bias towards happy faces is characterized by *γ* <0. Probabilities p_XYa_ and p_XYb_ are defined in Fig 2.

In the presented simulation, corrections of the responses were carried out only after acquiring a first estimate of the bias for each delay. A minimum of 25 trials and a minimum of at least 3 trials in each of the signal detection theory response categories (*P*_*CH*_ (*λ*), *P*_*CS*_ (*λ*), *P*_*IH*_ (*λ*), and *P*_*IS*_ (*λ*)) for the current delay *τ*, was required before the application of the correction procedure.

The flowchart (Fig 3) displays rules for the selection of a delay for the next trial, implemented through an algorithm that is based on corrected responses. After reaching a valid bias estimate, it is possible to correct an observer’s response by taking the initial bias value into account and using a new bias-free decision criterion set to *γ = 0*

In the AM, the average mask-delay of the last 50 trials was used as a threshold estimate after the preset number of trials had been reached. If, however, no predefined number of trials is defined after which the threshold procedure is stopped, a reasonable criterion for terminating the threshold estimation procedure is the reach of an asymptotic and stable threshold estimate within the last 50 trials. Using the two-down one-up rule, the probability of erroneously obtaining a stable threshold by chance is only 0.01 % for 30 trials and 0.0025 % for 50 trials and thus low enough to serve as an acceptable termination criterion for the AM threshold estimation procedure.

### Simulation Studies

#### Simulation of the virtual observer

To assess the comparative performance of the AM and MCS in the simulations, responses for the decision of a virtual observer for the presented stimuli are needed, respectively. The decision for a stimulus to be perceived by the virtual observer was simulated within the framework of SDT. The virtual observer’s detection competence was defined for ten delays Δ*t* (16.7 ms * k, with k ranging from 0 to 9). In order to cover the whole range of stimulus discriminability, spanning from most challenging to easy, d-prime for each delay was defined according to *d’* _Δ*t*_ = 12 Δ*t*.

The probabilistic decisions of how HF and SF were perceived, were based on Gaussian normal distributions centered at 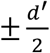 and with standard deviations *σ* of 1. Since the percentage of correct responses of 66.7%, i.e., correctly identified faces, corresponds to a *d’* of 0.861 the virtual observer’s detection threshold resulted in a mask delay of 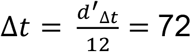. For the simulations, different observer’s biases (*γ*) were considered (0.0, 0.2, 0.5 standard deviations (*σ*) of the Gaussian distribution. Once the virtual observer’s detection competence, i.e., perceptual parameters, were defined, their performances for four different numbers of trials (100, 200, 500, and 1000 trials) were evaluated by simulations. To simulate the virtual observer’s decision, a z-score *r*_*s*_ was chosen from the normal distribution with a standard deviation of 1 and 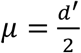 for HF and 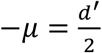 for SF. Depending on whether *r*_*s*_ exceeded the predefined decision criterion *γ: r*_*s*_ *≥ γ*,the virtual observer’s response was HF. Conversely, in case of *r*_*s*_ *< γ* the response was SF.

To evaluate the new method, a virtual observer’s performance for different numbers of trials and several levels of biases was simulated based on SDT and its threshold detection capabilities were estimated with and without bias correction, for both, the AM and MCS methods.

#### Simulation of AM for bias correction

For the simulation studies for the AM threshold estimation procedure, the virtual observer’s responses were fed into the AM and the stimulus for the next trial was selected accordingly. The sequence of stimulation was assumed to converge towards the preset threshold of 72 ms asymptotically. The results of the simulation for each set of parameters (level of biases and number of trials) were iterated for 1000 times and the corresponding thresholds were inferred (Fig. 4)

**Fig. 4:**
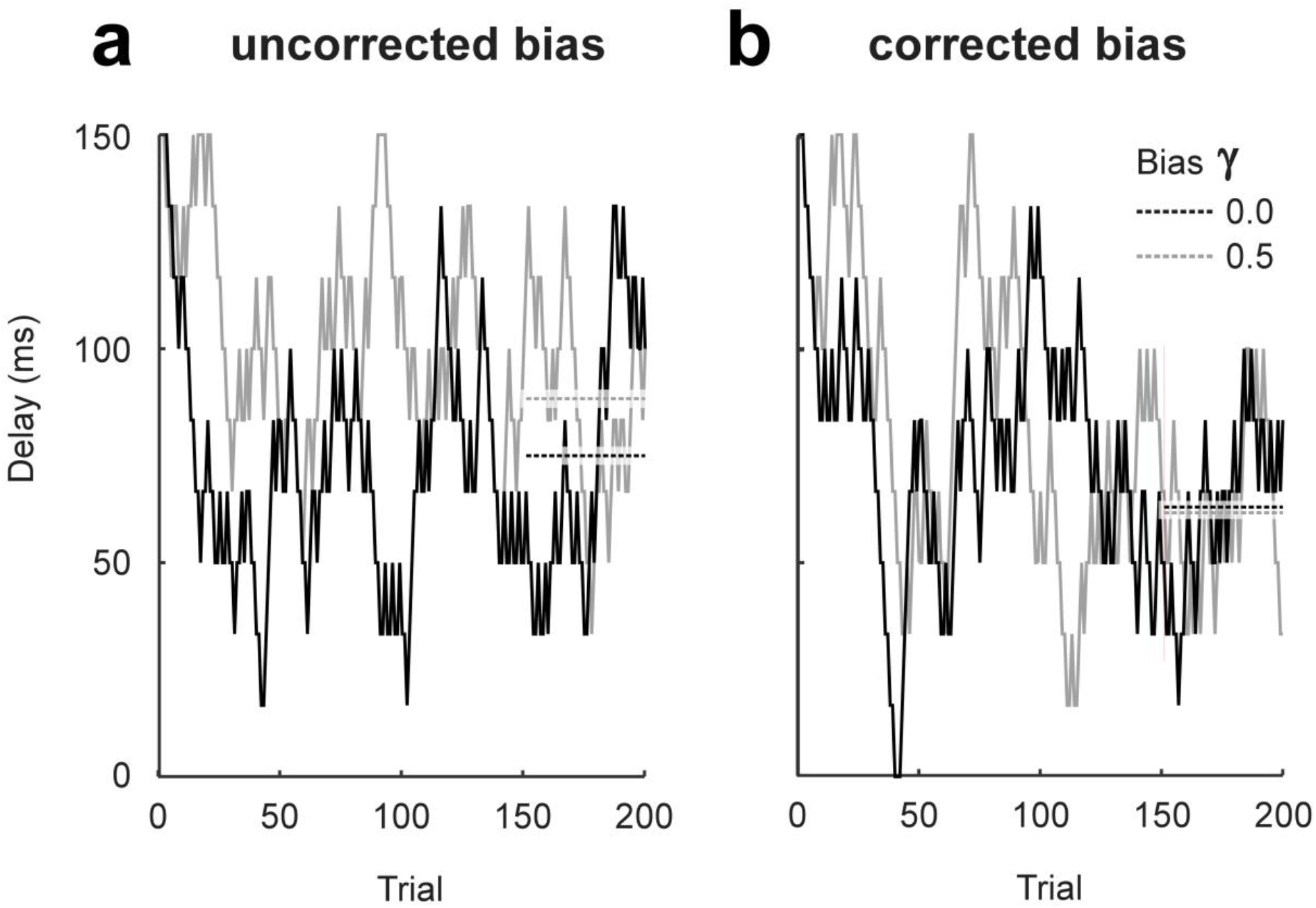
Simulation for the AM with an introduced bias of 0.0 (black) and 0.5 (grey); a: bias un-corrected threshold estimation, and b: bias-corrected threshold estimation. Threshold estimates were based on the average of mask delays for the last 50 trials. Averages are represented as horizontal dashed lines extending from trial 151 to 200.

#### Simulation of MCS

To contrast, the performance of the AM to a method for which bias correction had been already established, threshold estimates for AM, and MCS were compared using simulation studies. In the MCS approach, parameter settings for the virtual observer were identical to those used in the AM (see above). In the MCS, a psychometric function was fitted to the percentage of correct responses as a function of the ten different delays. Considering Fechner’s law (Fechner, 1860) of logarithmic relation between perceived and physical magnitudes of sensory input (Dehaene, 2003), the logarithms of all delays were calculated. To avoid the problem of obtaining a value of minus infinity for zero delays, 1 ms was added to all delays before the transformation. Afterward, a Weibull psychometric function was fitted to the percentage of correctly identified emotional face expressions:

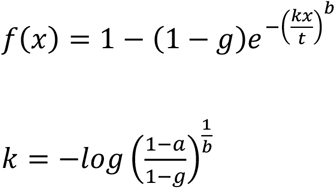

x = logarithmic transformation of delays

*g* = performance at the chance level: in our example set to 0.5

t = threshold

*a* = performance level defined as the threshold (0.667)

*b* = slope index of the psychometric function

The Weibull function asymptotically converges towards 50% for a delay of <1.0 ms and towards 100% for increasing delays (Weibull, 1938). The Weibull function fitted to the psychophysical data resulted in a threshold estimate for each observer, i.e., a delay for which a performance level of 66.7% was reached. The MCS threshold of 66.7% correct responses was chosen to comply with the two-down one-up procedure dependent threshold level of the AM.

#### Simulation of AM for time-varying sensitivities

Two possible scenarios were simulated to study the efficiency of the proposed adaptive procedure to track-changes/takes-into-account the adaptation of a virtual observer’s sensitivity across trials.

A) In first scenario, a successful perceptual training was simulated, assuming that the virtual observer’s sensitivity for detecting the emotion of the face stimuli improves linearly across 1000 trials. The apriori-sensitivity threshold was initialized at 100.0 ms and decreased by 0.033 ms/trial. After 500 trials, a sudden decline in the sensitivity is simulated by adding a threshold of 16.7 ms to explore the behavior of the algorithm for sudden changes. Thereafter, the threshold decreased at the same rate as at the beginning until it reached a value of 50.0 ms after 1000 trials.

B) In second scenario, which was inspired by slow variations in participants’ attention for threshold experiments, the performance of the virtual observer was simulated for randomly varying sensitivity. The random changes of the observer’s sensitivity were simulated by lowpass filtering white noise sampled at 1/trial such that fluctuations of threshold changes in the last 100 trials were suppressed. The virtual observers’ responses to the presented delays across trials were computed based on SDT (see paragraph ‘Simulation of the virtual observer’ above). For both the scenarios, the simulated observer’s perceptual biases were 0.0, 0.2, and 0.5, with and without bias correction.

#### Simulation of AM for a linearly changing bias

To investigate the performance of the algorithm for a linearly changing bias across trials, a linear change of the bias from 0.0 to -1.0 and from 0.0 to +1.0 across 1000 trials with steps of 0.001, was simulated. The threshold estimation used a two-down one-up procedure resulted in 66.7 %. The results were presented for runs with and without bias correction.

## Results

The simulation results suggest that when the number of trials is less than 200, the MCS method to estimate threshold fails to reliably fit a Weibull function, both, for corrected and uncorrected bias thresholds. On the other hand, AM is not constrained by this situation. As expected, in the bias uncorrected AM procedure, the estimated thresholds are independent of the number of trials. Moreover, the bias-corrected AM managed to approach the threshold that was preset in the simulation. Since the threshold of a single individual obtained with one method (corrected or uncorrected AM), falls within the 95 % confidence interval (± 2 standard deviations) of the other method (uncorrected and corrected AM), individual threshold values do not differ significantly between corrected AM and uncorrected AM. However, comparing the mean of a group of *N* subjects would reduce the confidence interval by a factor of 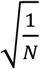 of the standard deviation. Threshold estimates are comparatively lower in MCS than in the AM across the number of trials and for different preset biases. Furthermore, threshold estimates using the AM method are independent of the number of trials required for the reliable estimate as indicated by the constant standard deviation across the number of trials for a different level of simulated perceptual bias. In contrast, threshold estimates for the MCS method rely variably on the number of the trials as suggested by the decreasing trend in the standard deviation across trials and levels of preset bias. (Fig. 5). This finding further suggests that the threshold estimates using AM method do not require criteria for a minimum number of trials whereas the MCS method does have a requirement of minimum trials as it failed to fit the Weibull function reliably for the condition with less than 200 trials. To quantitatively assess the performance of the algorithm, we studied the threshold estimated with or without bias correction for both AM and MCS methods against the predefined threshold for different trial numbers and at different simulated bias levels against the predefined inputs in the simulations. The accuracy of the bias estimate, i.e., its deviation from the simulated preset bias value, and its precision, reflected by the standard deviation (the lower the standard deviation, the higher the precision), grew with increasing trial numbers for both methods. The assessment of performance suggests that the number of trials required to achieve the preset bias for both methods is comparable, whereas the precision of the estimate is comparatively better for the AM as suggested by the small standard deviation across trial numbers and across different levels of introduced bias (Fig. 6).

**Fig. 5:**
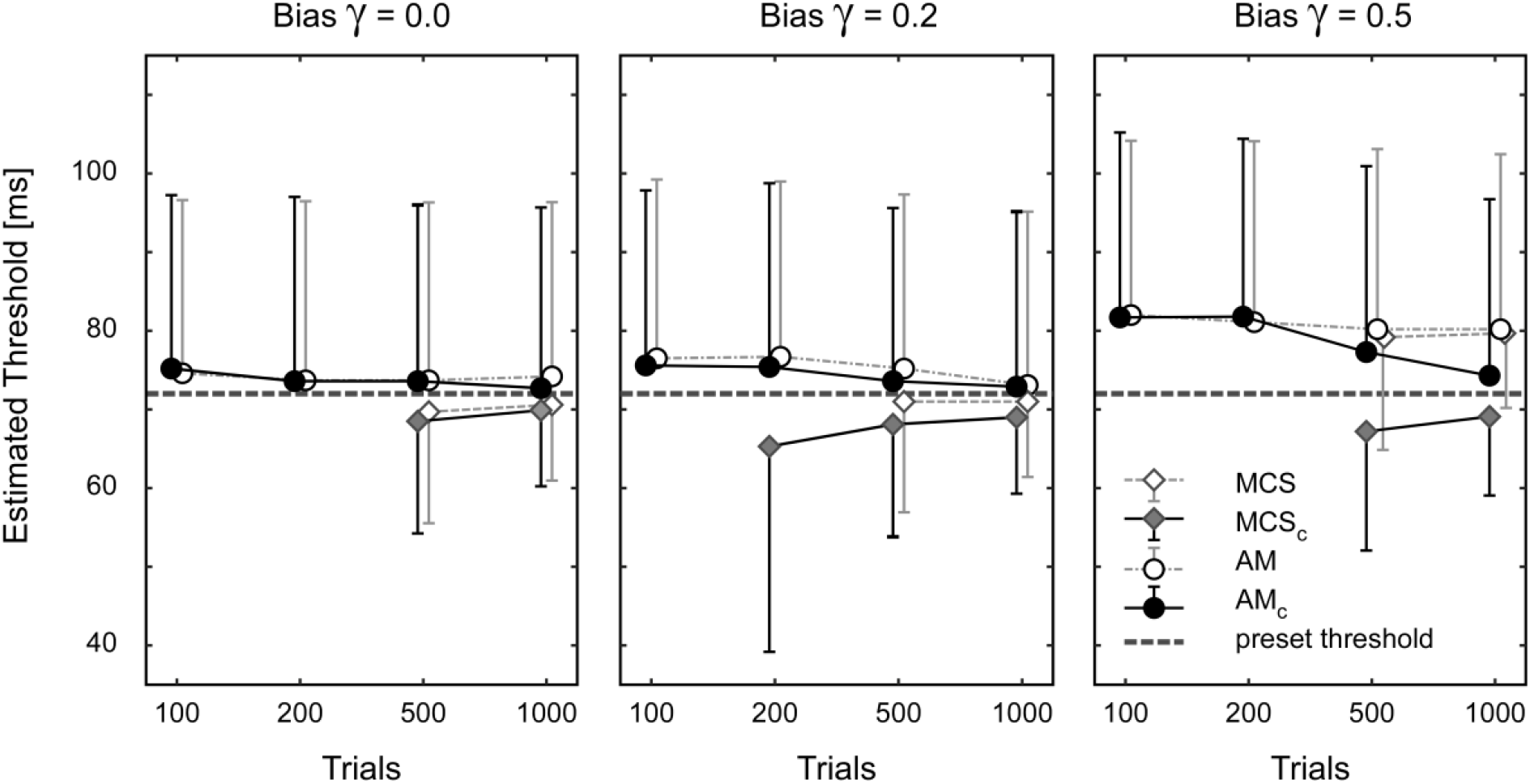
Threshold estimates of the AM and MCS with and without bias correction for a bias of 0.0, 0.2, and 0.5 (left, middle and right part) and trial numbers of 100, 200, 500, 1000 (abscissa). For each condition, mean and standard deviations across 1000 simulations are presented (to improve visibility error bars are plotted only in one direction). Missing results for 100 and 200 trials are due to the failure of reliably fitting a sigmoid Weibull function and thus being unable to estimate a threshold value. Bias correction is based on the bias estimate, derived from the observer’s previous responses. The bias values are related to the width of the normal distribution, describing the variability of the stimulus perception.

**Fig. 6:**
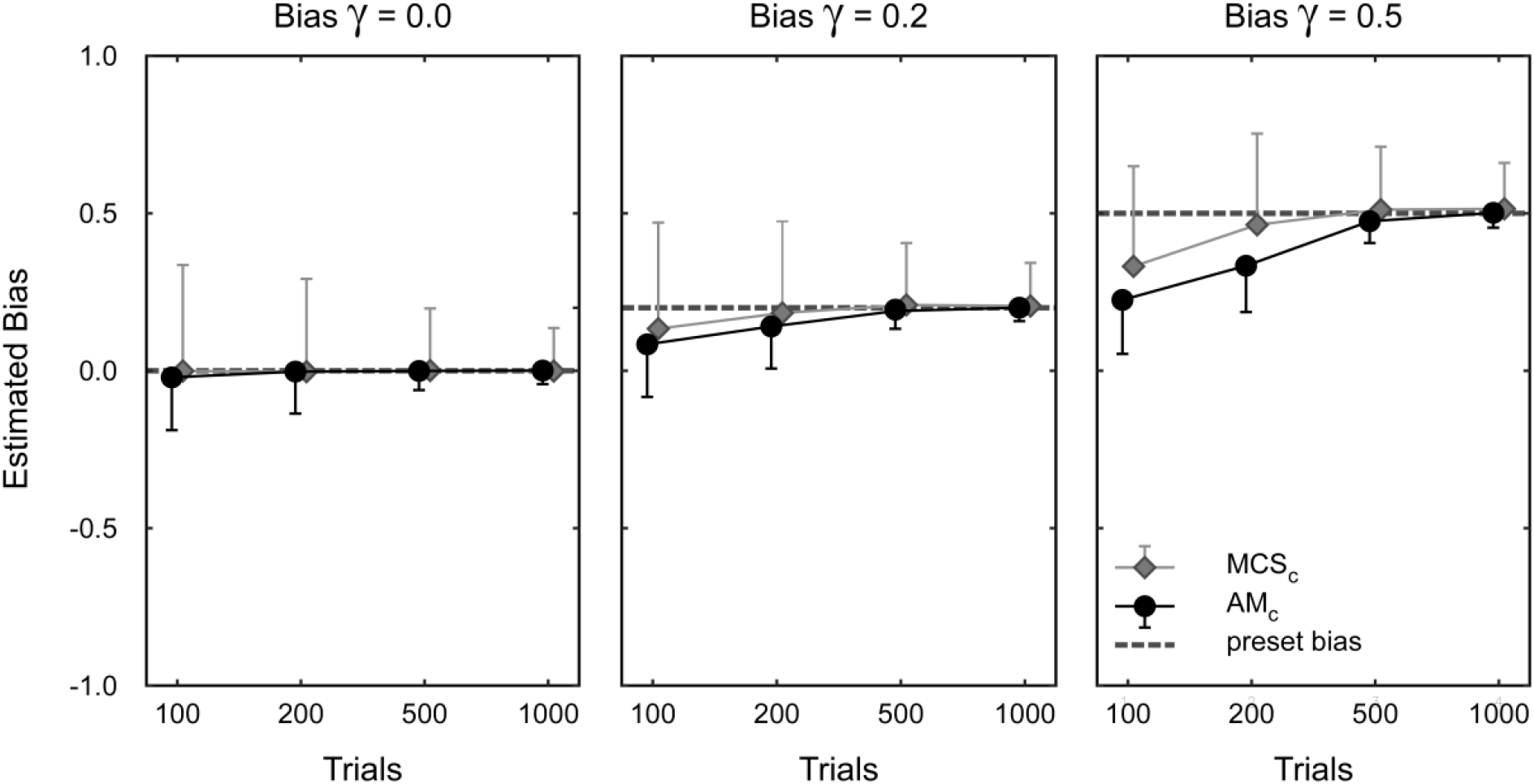
Estimated biases values determined by the AM and the MCS with bias correction. Mean bias estimates 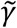 and standard deviations 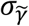 across 1000 simulations are presented for simulated decision criteria *γ* of 0.0, 0.2, and 0.5 and for trial numbers of 100, 200, 500, and 1000 (abscissa).

To study the performance of the algorithm considering the change of the perceptual sensitivity across trials, we simulated a linear change in the sensitivity profile. Our results as shown in figure 7 demonstrate that our algorithm for AM is capable of tracking the participant’s threshold continuously with changing the sensitivity, yet with a lag of about 50 trials for both the AM method. Also, the AM estimated threshold follows the preset threshold of the virtual observer in a smoothened way, due to the hysteresis of the AM procedure. Due to the steady change of the virtual observer’s sensitivity and the delay of the estimation procedure, it is clear that the threshold estimate cannot fully converge to the virtual observer’s current threshold. For a randomly varying sensitivity, the AM threshold procedure tracks the fluctuation of the preset sensitivity as long as their time constants are well above the time interval across which the threshold parameters are averaged. Interestingly, for the variable sensitivity of the virtual observer the latter approach yielded also more precise threshold estimates than the two-down one-up approach. In the simulations without any bias correction, the accuracy of the threshold estimate was best for zero bias and worst for a bias of 0.5. The bias correction showed a good outcome especially for the virtual observer’s randomly varying sensitivity. Also, the bias correction mechanism is switched on relatively early for intensities around the threshold (step-like lines for the different biases in figure 7 panel b) and d), it requires a large number of trials for the procedure to become effective for mask delays further away from the threshold.

**Fig 7:**
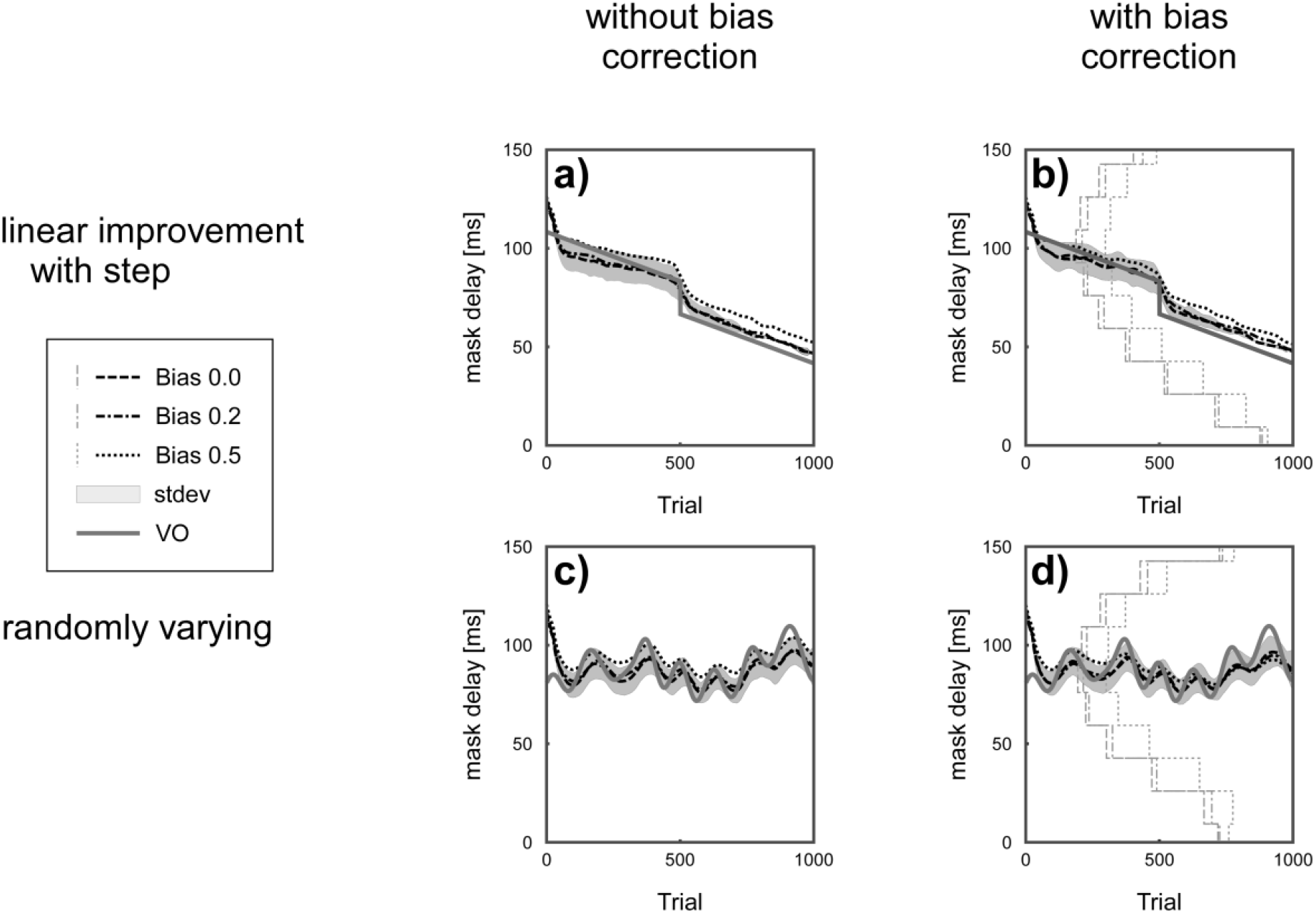
Simulation of the performance of the AM for a virtual observer’s changing sensitivity across trials. In scenario 1 (panel a) and b)) the virtual observer’s sensitivity, i.e., threshold, improves linearly from 100.0 ms to 83.3 ms from trial 1 to trial 500. At trial 501 a sudden change of the improvement (16.7 ms) was simulated to demonstrate the behavior of the AM to fast changes. From trial 501 onwards to trial 1000 the virtual observer’s threshold further decreased again linearly resulting in a final threshold of 50.0 ms in trial 1000. In scenario 2 (panel c) and d)) a randomly changing virtual observer’s threshold was simulated. In both scenarios’ biases of 0.0 (dashed lines), 0.2 (dashed-dotted lines), and 0.5 (dotted lines) were simulated. The AM procedure was run with a two-down one-up procedure resulting in a threshold performance of 66.7 % correct responses. The threshold estimation was done without (a) and c)) and with bias correction (b) and d)). The standard deviation of the threshold estimates for 1000 repetitions for the zero-bias simulation is shown as a grey area. The step-like grey lines indicate the average trial number at which the bias correction was switched on. The trial at which the correction becomes active varies for the different mask delays of the stimuli and for the virtual observer’s bias (0.0: solid lines, 0.2: dashed-dotted lines, and 0.5: dotted lines). Since the estimated threshold follows the simulated threshold only after a delay, the estimated threshold is shifted by around 50 trials to the right. Grey dashed lines for the increasing bias and grey dotted lines of the decreasing bias. In case of no bias correction there is a constant offset of the estimated threshold with respect to the preset threshold of the virtual observer. The bias correction works well for randomly varying thresholds of the virtual observer yet fails for the steady improvement of the virtual observer’s sensitivity.

To investigate how the bias correction procedure deals with systematically changing biases over trials, linearly changing biases were simulated for a preset threshold delay of 72 ms. While in condition one the bias was increasing linearly from 0 to 1 across trials 0 to 1000, it was decreasing from 0 to -1 in condition two. The effects of decreasing and increasing biases on the virtual observer’s threshold did not differ (Fig 7). Comparing the estimated thresholds with and without bias correction, the threshold deviated less from the preset threshold of 72 ms if the correction was activated. However, the bias correction was unable to fully eliminate the simulated linear bias (Fig. 8).

**Fig. 8.**
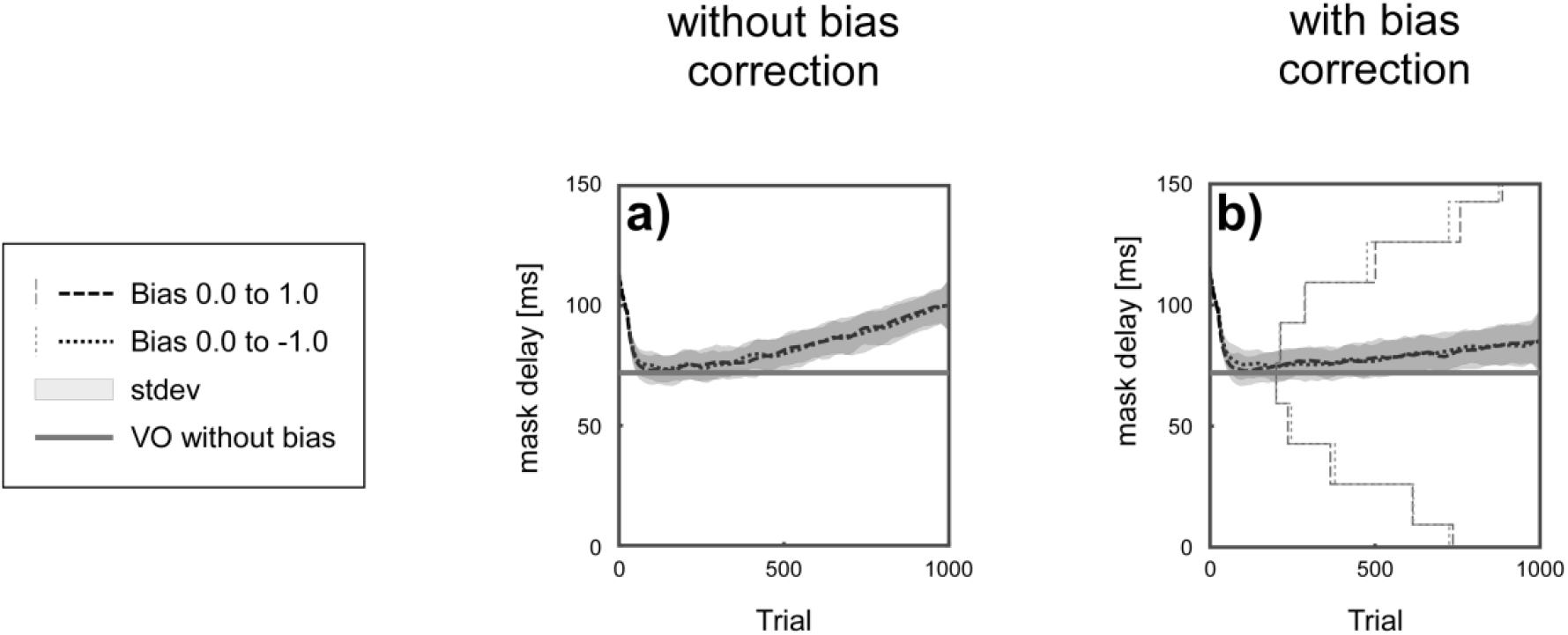
Simulation of a linearly changing bias across trials. The solid grey horizontal line represents the preset performance of the virtual observer (VO) without any bias. Thresholds estimates for a steady bias increase from 0.0 to 1.0 and a decrease from 0.0 to -1.0 are depicted in dashed and dotted lines, yet they are indistinguishable. Grey areas indicate the standard deviation of the threshold. Step like lines indicate the trials at which the bias correction became active (grey dashed lines for the increasing bias and grey dotted lines of the decreasing bias). In contrast to panel a) bias correction was activated in panel b). Applying the two-down one-up rule resulted in a threshold performance of 66.7 %.

## Discussion

In the current piece of work, we proposed a novel approach for assessing and correcting the observer’s bias in an adaptive threshold procedure (AM) for estimating the perceptual threshold. We performed various simulations studies with a virtual experiment in which the virtual observer’s threshold in a backward masking task was estimated using the two-down one-up rule. Simulations of the virtual observer’s sensory performance investigated various factors affecting the perceptual threshold, such as a bias for a specific stimulus class, and linearly changing perceptual biases across trials. Moreover, effects of various levels of time-dependent sensitivity profiles were studied with scenarios such as linearly and randomly changing sensitivities across trials using the novel AM procedure. Using simulation studies, we have explored the performance of the algorithm and compared the obtained results against the standard procedure of the threshold estimation using MCS.

Previously suggested threshold estimation procedures that considered sensitivity and bias, have used Bayes’ theory – with an apriori distribution of probabilities for threshold parameters, including sensitivity and bias, and estimates their posterior probabilities based on the sampled data (Lesmes et al., 2015). In the method proposed by van Dam and Ernst et. al, the observer’s bias is assessed through a set of Kalman Filters (Rohde, van Dam, & Ernst, 2016). In contrast to these class of methods, the here proposed procedure does not rely on any prior assumption and any apriori knowledge.

Using simulations, the performance of the bias-corrected threshold estimation by an adaptive method is compared to the threshold estimates of the method of constant stimuli. To show whether the limitations we encounter are specific to the proposed procedure or are a general issue of threshold estimation procedures, employing bias correction, we used a standard MCS method as a reference that excluded any response-dependent sampling near the threshold.

Results indicate that both, AM and MCS, were able to estimate sensitivity and bias correctly, in case of a sufficiently large number of trials (>200 or > 500). In case of a trial number ranging between 100 and 200, the MCS method however fails to reliably estimate the threshold even without any bias correction. On the contrary, the AM procedure is capable of estimation the perceptual threshold with and without bias correction independently from the number of trials (Leek, 2001; Lesmes et al., 2015; McKee et al., 1985; Treutwein, 1995; Watson & Fitzhugh, 1990).

In the MCS method, a psychometric function is fitted to the percentage of correct responses as a function of the delay. If responses for single delays are too noisy as in the case of few trials, then the fitting of the psychometric function might be corrupted and will eventually fail. The failure of the MCS for low trial numbers, in the chosen example of the backward masking paradigm, results from the necessity of this approach to equate the total number of trials for different mask delays (Leek, 2001; Treutwein, 1995; Watson & Fitzhugh, 1990), leaving only a fraction of trials for a single delay. In contrast, in AMs, the delays in individual trials are mostly concentrated around the threshold delays, and thus a larger number of trials is available for calculating the threshold estimates. MCS versions that sample the observer’s responses close to the threshold with higher numbers of trials are certainly less affected by this problem, yet involve the response-dependent selection of the sampling region – a characteristic feature of AM.

Results of the bias-uncorrected AM reveal that the threshold estimate does not vary much with the number of trials, at least as long as the threshold remains constant. Assuming that the delays used for simulations in the AM converge towards the threshold quickly, and using the averaged delay of the last 50 trials in each run, the threshold delays should be independent of the number of trials. This is especially true for trial numbers larger than 200 (Fig. 5). Since in the presented example there were only 10 steps of delays, the lowest delay (0.0 ms) could be reached within 20 trials using the two-down one-up rule, even if the procedure starts at the maximum delay (150 ms).

With increasing numbers of trials, the bias-corrected AM yielded threshold estimates approaching the value preset in the simulation. In the AM method, bias-corrected estimates of the threshold rely on the bias-corrected responses of the observer, that determine the stimulus delay for the next trial according to an *m*-down *n*-up rule requiring “*m”* correct response trials and “*n”* incorrect response trials before decreasing or increasing the delay in the upcoming trial, respectively. A first, estimate of the bias is generated during the initial trials. Afterward, when the bias estimate is available, the algorithm for bias correction comes into play. Assuming a constant bias across the experiment, the bias estimate is constantly updated and thus becomes increasingly more reliable. However, this strategy implies that a sufficiently high number of trials had been sampled to get a good estimate of the bias. The simulations show that it takes around 200-500 trials in the MCS and the AM, respectively.

The convergence to the bias-corrected threshold becomes slower with increasing biases. The reason for this relation is the fact that the tails of the probability density function, defined in the SDT, become smaller with more extreme bias values and thus increasingly difficult to estimate. For instance, in case of a decision criterion shifted towards ‘sad’ favoring more ‘happy face’ responses (see Fig. 2), the slow convergence is due to the low probability of obtaining an incorrect response when presenting a happy face (Fig. 2a). Similarly, in case of a shift of the decision criterion towards ‘happy’, the probability for obtaining an incorrect judgment when presenting a sad face will be low (Fig. 2d). With the low probabilities of responses, the time that is needed to reach a sufficiently high number of trials to reliably estimate the bias, increases drastically. However – as shown in the simulations of time-varying thresholds of the virtual observer for stimulus parameters close to the threshold – more trials are available and thus the procedure for bias correction is switched on earlier than for stimuli further away from the threshold.

In summary, for both the AM and the MCS, bias correction improves the threshold estimation accuracy, especially in case of strong biases. However, an effective bias correction algorithm requires a relatively large number of trials (see: S1a-b). Since for stronger biases, the required number of trials to obtain a stable estimate is large for both methods, and therefore certain advantages of the AM over the MCS (fewer trials needed to estimate the threshold) might get lost.

If the mean threshold of a larger group of individuals is studied, differences between methods will become more evident. Since the standard error of the mean will decrease by the square root of N, with N being the number of individuals, the minimum group size, for which the application of bias correction is beneficial, can be inferred.

With a sufficient number of subjects or repetitions, threshold estimates could be consistently higher for the AM as compared to the MCS in settings similar to our virtual experiment: The systematic higher threshold for the AM might be a result of the lower limit of Δ*t* = 0 the mask delay. In the case of two correctly perceived emotional face expressions presented with a mask delay of 0 ms, the delay should be further decreased according to the two-down one-up rule. However, reducing the delay below 0 ms is not feasible in our experiment, and thus, delays for low thresholds in the AM will be slightly overestimated. The reliability of threshold estimates that are indexed by the standard deviation across simulation repetitions, is constant for the AM, independently of the number of trials and bias levels. In contrast to MCS, the standard deviations of different threshold levels are rather high, rendering the AM less reliable. With the two-down one-up rule applied here, only a few of the most recent trials determine the variation of the stimulus delay in the upcoming trials, making the approach very sensitive to noise. An alternative rule that considers responses of a higher number of “most recent” trials, reduces the effects of noise and, as has been shown in our simulation, will enable the estimation of threshold levels for performance levels higher than 66.7%.

Results for the simulation of a linear changing bias showed similar effects on the virtual observer’s threshold for decreasing and increasing biases (Fig. 8). This result is to be expected because flipping the sign of the decision criterion does not affect the overall correct responses, which is the sum of the numbers for hits and correct rejections. The bias-corrected threshold estimate deviated less from the preset threshold of 72 ms compared with the uncorrected one. However, the correction procedure was unable to fully correct the simulated bias changing across trials. This finding can be explained by the fact that the bias estimate used for the correction of the bias in a current trial is based on the observer’s responses in all previous trials and therefore most likely not fitting the situation in the current trial. In case of a constantly increasing bias across trials, the bias in a certain trial is always underestimated and thus the correction is incomplete.

In general, a better threshold and bias estimate are expected from the adaptive method because there are more trials near the threshold. However, assuming a systematic relation between mask delay and discrimination performance in the psychometric function, like we did in our simulations, fitting a steady psychometric function to all data points in the MCS, might compensate for the higher error near the threshold.

While in this article, bias was regarded as a nuisance parameter that masks subjects’ sensory capacities (Witte, Kober, Ninaus, Neuper, & Wood, 2013), it is important to underline that in other studies, changes in bias are the parameters of interest. For instance, studying the perception of emotional stimuli in psychiatric diseases, such as depression (Bourne & Vladeanu, 2013), schizophrenia (Gooding & Tallent, 2002) or autism (Ashwin, Wheelwright, & Baron-Cohen, 2005; Taylor, Workman, & Yeomans, 2012) or in healthy subjects (Kajal, 2018; Kajal et al., 2017; Kajal et al., 2018), the observed higher or lower thresholds might result from a shift in the sensory bias rather than from altered sensitivity. Since the proposed method differentiates sensitivity and bias, deriving estimates for both, the method might have a wide range of applications in psychotherapy, in which would be interesting to modulate these parameters independently.

## Conclusion

In this study, a new adaptive threshold estimation procedure was introduced, which can correct an observer’s bias reliably. The performance of the new procedure was simulated and compared to other approaches. Furthermore, the study provides insight into the performance of classical threshold estimation procedures with and without bias correction and discloses limitations of the procedures in this context.

Comparable number of trials are required for both AM and MCS procedure for a reliable threshold estimate. To minimize the effects of observers’ bias, either time-consuming correction procedures can be applied, or experiments should be designed more carefully minimizing effects of biases. In general, MCS indicates better reliability than AM, yet, at the cost of a large number of trials. In contrast, AM with bias correction is especially beneficial in case of low bias values as it requires less numbers of trials for reliable threshold estimates. The AM with bias correction are the method of choice in experiments where the observer’s threshold is dynamically fluctuating. Experiments in which sensory performance is sought to change, the continuous adaptation of sensory stimulus parameters to the current perceptual threshold allows maintaining tasks demands constant across the whole experiment. Although the methodological framework presented in this study leaves space for further improvements, the new approach reveals a promising potential with a relevant impact on psychophysics, behavioral learning, and neurofeedback training.

## Acknowledgments

This project was realized with the support of the Werner Reichardt Centre for Integrative Neuroscience (CIN) at the University of Tübingen. The CIN is an Excellence Cluster funded by the Deutsche Forschungsgemeinschaft (DFG) within the framework of the Excellence Initiative (EXC 307). Furthermore, the research was supported by the DFG-grant BR 1689/9-1; Comisión Nacional de Investigación Científica y Tecnológica de Chile (Conicyt) through Fondo Nacional de Desarrollo Científico y Tecnológico, Fondecyt Regular (projects n◦ 1171313 and n◦ 117132) and CONICYT PIA /Anillo de Investigación en Ciencia y Tecnología ACT172121, The Cockrell School of Engineering, The University of Texas and School of Engineering, as well as the Pontificia Universidad Católica de Chile.

We would also like to thank Mr. Jürgen Dax for coding and testing the stimulus presentation software, and Dr. Thomas Hess for proof-reading the manuscript.

## Data and code availability statement

The data that support the findings of this study are available from the author on request.

